# Resolving the molecular fingerprint of the distal carboxy tail in modulating Ca_V_1 calcium dependent inactivation

**DOI:** 10.1101/2021.01.06.425618

**Authors:** Lingjie Sang, Daiana C. O. Vieira, David T. Yue, Manu Ben-Johny, Ivy E. Dick

## Abstract

Ca^2+^/calmodulin-dependent inactivation (CDI) of Ca_V_ channels is a critical regulatory process required for tuning the kinetics of Ca^2+^ entry for different cell types and physiologic responses. Calmodulin (CaM) resides on the IQ domain of the Ca_V_ carboxy-tail, such that Ca^2+^ binding initiates a reduction in channel open probability, manifesting as CDI. This regulatory process exerts a significant impact on Ca^2+^ entry and is tailored by alternative splicing. Ca_V_1.3 and Ca_V_1.4 feature a long-carboxy-tail splice variant that modulates CDI through a competitive mechanism. In these channels, the distal-carboxy-tail (DCT) harbors an inhibitor of CDI (ICDI) module that competitively displaces CaM from the IQ domain, thereby diminishing CDI. While this overall mechanism is now well-described, the detailed interaction loci for ICDI binding to the IQ domain is yet to be elucidated. Here, we perform alanine-scanning mutagenesis of the IQ and ICDI domains and evaluate the contribution of neighboring regions. We identify multiple critical residues within the IQ domain, ICDI and the nearby A region of the channel, which are required for high affinity IQ/ICDI binding. Importantly, disruption of this interaction commensurately diminishes ICDI function, as seen by the re-emergence of CDI in mutant channels. Furthermore, analysis of the homologous ICDI region of Ca_V_1.2 reveals a selective effect of this channel region on Ca_V_1.3 channels, implicating a cross-channel modulatory scheme in cells expressing both channel subtypes. In all, these findings provide new insights into a molecular rheostat that fine tunes Ca^2+^ entry and supports normal neuronal and cardiac function.

## Introduction

L-type voltage-gated calcium channels (Ca_V_1.1-1.4) are an important conduit for extracellular Ca^2+^ entry into many excitable cells including cardiac myocytes, neurons, smooth muscle and skeletal muscle (1–4). As such, these channels are subject to rich and powerful modes of feedback regulation (5–7). In particular, Ca^2+^ dependent inactivation (CDI) of L-type channels is a crucial negative feedback mechanism that reshapes the electrical properties of neurons and cardiac myocytes and protects cells from Ca^2+^ overload (8–10). CDI is driven by the ubiquitous Ca^2+^ sensing molecule, calmodulin (CaM) (10,11). Under basal Ca^2+^ conditions, Ca^2+^-free CaM (apoCaM) binds to the carboxy-terminal IQ domain of the channel and enhances channel openings (12). Upon elevation of intracellular Ca^2+^, the ‘resident’ CaM repositions on the channel, interacting with Ca^2+^/CaM binding sites located on the channel amino- and proximal carboxy-termini (13,14). This conformational change antagonizes the initial upregulation in channel open probability, which manifests as CDI. Not surprisingly, CDI of L-type channels has emerged as a key physiological process to limit excess Ca^2+^ influx during repetitive or sustained depolarization, and disruption of this feedback in the cardiac myocytes may lead to lethal cardiac arrhythmias (15,16). This stereotypic behavior, however, diverges in multiple physiological settings where strong CDI of L-type channels is curtailed, thus permitting Ca^2+^ channels to faithfully respond to a tonic stimulus. For example, in photoreceptors and bipolar cells, endogenous Ca_V_1.4 exhibit minimal CDI, thereby allowing sustained Ca^2+^ influx and slow, graded changes in the membrane potential necessary for tonic glutamate release and normal vision (17,18). Similarly, Ca_V_1.3 channels in the inner hair cells also lack CDI (19). Beyond these, the basal strength of Ca_V_1.2 and Ca_V_1.3 CDI vary in different neuronal subtypes in the central nervous system, suggesting a sophisticated scheme of Ca_V_ channel feedback ripe with physiological insights (20).

The molecular mechanisms that fine-tune L-type channel CDI are two-fold and have been of long-standing interest. One scheme involves channel-interacting proteins such as calmodulin-like Ca^2+^-binding proteins (CaBP1-4) (18,19,21,22) and SH3 and cysteine rich domain containing proteins (stac1-3) (23–26) that suppress CDI utilizing an allosteric or mixed-allosteric mechanism. In contrast, Ca_V_1.3 and Ca_V_1.4 channels may intrinsically disable CDI via an alternatively-spliced specialized CDI-inhibiting module (ICDI) within the distal carboxy-tail of the channel (20,27–33). The latter form of regulation is complex and bears important biological consequences. First, splice-inclusion of the distal carboxy-tail occurs in a cell-type dependent manner. For instance, alternative splicing of Ca_V_1.3 results in variable inclusion of the ICDI domain in distinct regions of the brain and in the sinoatrial node, enabling precise tuning of CDI in these cell types (27,31,34,35). Second, Ca_V_1.2 appears to harbor a highly homologous ICDI region (33), yet CDI for this channel is known to be robust, both when evaluated as full length channels in heterologous expression system, as well as in primary cells where the C-tail containing ICDI is believed to be cleaved off the channel (9–11,33). As such, the function of the ICDI module within Ca_V_1.2 channels remains unclear. Third, the inhibition of CDI by ICDI is the result of competitive binding by apoCaM *versus* ICDI with the channel IQ domain (20,31). In addition to diminishing CDI, the displacement of apoCaM results in a dramatic decrease in baseline channel open probability (12). Fourth, adding to the richness the modulatory role of ICDI, RNA editing and/or fluctuations in cytosolic CaM concentrations can tune the extent of this competition, enabling different degrees of CDI tailored to specific cell types or physiologic states (12,36). Importantly, pathologic changes to this system may be linked to altered CaM concentrations in Parkinson’s disease and heart failure (37,38), and mutations within the ICDI of Ca_V_1.4 channels are known to be causative of congenital stationary night blindness (32,39,40). Thus, the modulation of CDI by ICDI stands as a critical and robust mechanism for adapting channel regulation to select cell types and conditions. Moreover, as the number of known pathogenic mutations within LTCCs continues to grow, the ability to map these mutations to a locus with known mechanistic impact would enable rapid insight into the pathogenesis of LTCC channelopathies.

Although the overall competitive nature of ICDI regulation of L-type channels is now well-established (20,31), the precise binding interfaces involved in this regulation are yet to be identified. This gap in understanding is critical as mutations in the carboxy-tail of LTCCs result in neurological disease (32,39,40). Furthermore, a residue-level analysis may shed light upon structural differences between the ICDI domain of Ca_V_1.3 and the homologous segment of Ca_V_1.2 that engender differential functional regulation. To characterize the landscape of the IQ/ICDI interaction of L-type channels, here we undertook systematic alanine scanning mutagenesis of both IQ and ICDI domains. Through live-cell FRET two-hybrid binding assays and electrophysiological analysis, we identified several novel hotspots on both IQ and ICDI segments that mediate a high-affinity interaction and are functionally relevant for CDI inhibition. Systematic analysis of these mutations revealed a strong inverse correlation between the strength of CDI and the binding affinity of the ICDI domain for the IQ segment, as predicted for a competitive inhibitor (13,14,20). Thus, we have identified residues which alter binding in a functionally relevant manner. Moreover, similar critical residues were identified in adjacent regions, defining a comprehensive interface map of the IQ/ICDI interaction. Finally, extending our analysis to Ca_V_1.2 channels, we found that the ICDI module binds to the Ca_V_1.2 IQ domain with a reduced affinity, and that this binding is insufficient to cause more than a nominal change in the CDI of full length channels. However, the ICDI from Ca_V_1.2 is capable of binding the Ca_V_1.3 IQ region with high affinity, resulting in a much larger decrease in CDI of these channels. Given the propensity of the C-tail of Ca_V_1.2 to exist as a separate peptide within myocytes and neurons(27,33,41,42), these findings raise the prospect of a cross-channel feedback scheme in some cell types. Overall, these results elucidate the detailed binding interface between components of the carboxy-tail of L-type Ca^2+^ channels, lending new insight into normal and pathologic channel regulation.

## Results

### Identification of critical residues within the IQ domain necessary for ICDI

To identify key residues that support a high-affinity IQ/ICDI interaction, we undertook systematic alanine substitution of the IQ domain and evaluated both the relative binding affinity and the strength of ICDI mediated inhibition of CaM regulation. Importantly, the ICDI domains of both Ca_V_1.3 and Ca_V_1.4 are highly homologous, and have been shown to interact with IQ domains in like manner evoking similar functional effects (20,27,29). Even so, the ICDI domain from Ca_V_1.4 (ICDI_1.4_) has a greater binding affinity for the IQ domains of both Ca_V_1.3 and Ca_V_1.4, with FRET binding assays yielding more robust measurements with enhanced signal to noise ratio as compared to ICDI_1.3_ (43). We therefore focus on this canonical ICDI motif for our studies. However, robust expression of the holo-Ca_V_1.4 channel in recombinant systems is notoriously challenging, largely due to their diminutive open probability (12,44). We therefore chose to explore the interaction between the IQ domain of Ca_V_1.3 channels (IQ_1.3_), and ICDI_1.4_. To this end, we utilized a chimeric channel in which the DCT of Ca_V_1.4 is spliced onto the backbone of Ca_V_1.3 (Ca_V_1.3_Δ/DCT1.4_) (**Fig. 1A**), which has previously proven useful in dissecting the mechanisms underlying ICDI modulation of the channel (12,20). This chimera furnishes a strong IQ/ICDI interaction coupled with a robust functional readout, enabling quantitative analysis.

**Figure 1.**
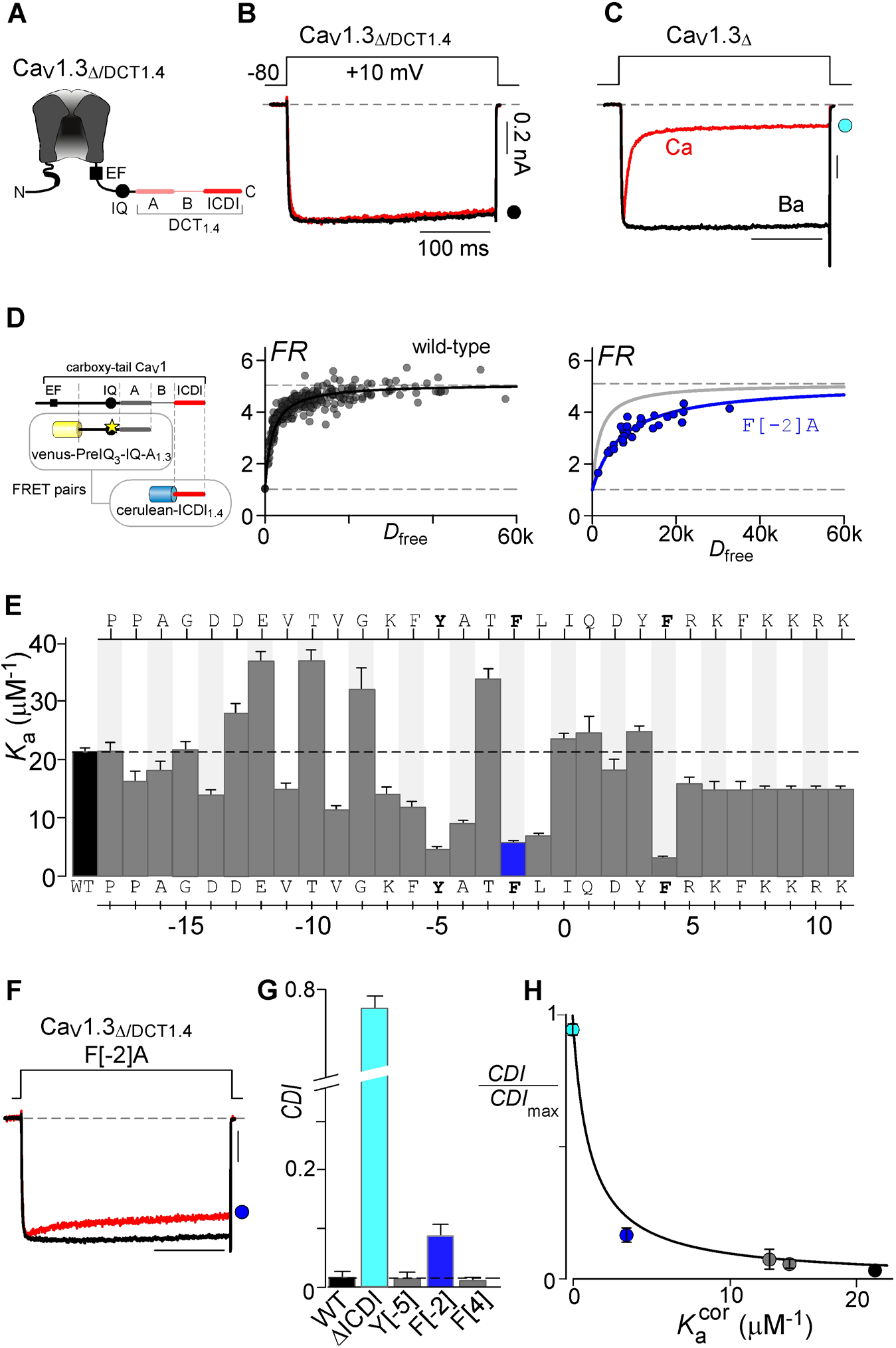
Identifying the hotspots on the IQ region required for ICDI binding and function. (**A**) Cartoon depicting the chimeric Ca_V_1.3_Δ/DCT1.4_ channel. (**B**) Exemplar whole cell recording showing minimal inactivation in response to a 10mv step depolarization in Ba^2+^ (black) or Ca^2+^ (red) for Ca_V_1.3_Δ/DCT1.4_ demonstrating the function of ICDI_1.4_. Ba^2+^ current is scaled to enable comparison of the kinetics of the two traces. Scale bar refers to the Ca^2+^ trace here and throughout. (**C**) Exemplar whole cell recording of the truncated Ca_V_1.3 channel, lacking the DCT containing ICDI. Robust CDI is seen as the strong decay of the Ca^2+^ current (red) as compared to the Ba^2+^ current. (**D**) FRET two-hybrid assay of IQ/ICDI interaction. FRET binding partners are displayed on the left, with star indicating the locus of alanine mutations. Strong binding was measured for wild-type PreIQ_3_-IQ-A_1.3_ with ICDI_1.4_ (black), while the mutation F[-2]A (blue) decreased the binding affinity compared to WT (gray). (**E**) Summary of *K*_a_ values for mutant venus-PreIQ_3_-IQ-A_1.3_ *versus* Cerulean-ICDI_1.4_ peptides measured with FRET two-hybrid assay as in panel D. Alanine was systematically substituted into the IQ region of the venus-PreIQ_3_-IQ-A_1.3_ peptide, with the identity of the amino acid displayed on the top and bottom of the bar graph such that the canonical ‘I’ of the IQ region is given position 0. The dashed line indicates the WT value and the blue bar corresponds to F[-2]A displayed in panel D. Data are displayed as mean ± SEM. (**F**) Exemplar patch clamp data corresponding to F[-2]A in Ca_V_1.3_Δ/DCT1.4_ demonstrating a partial recovery of CDI due to the mutation. (**G**) Average CDI values for each mutation, colors correspond to the data in other panels. (**H**) CDI and binding data for the mutations is well fit by equation 2, validating the competitive mechanism. Colored circles correspond to the colored data in the figure panels.

To begin, we confirm the functional impact of ICDI in our chimeric channel by evaluating the extent of CDI in HEK293 cells. Indeed, CDI is entirely abolished in Ca_V_1.3_Δ1626/DCT1.4_, as seen by the identical Ba^2+^ and Ca^2+^ current decay in response to a depolarizing pulse (**Fig. 1B**). However, removal of DCT_1.4_ restores robust CDI, as seen by the rapid decay of the Ca^2+^ current (**Fig. 1C,** red). In contrast, when Ba^2+^ (which binds poorly to CaM) is used as the charge carrier, there is minimal inactivation (**Fig. 1C**, black). We therefore define the extent of CDI as the ratio of Ca^2+^ current remaining after 300 ms of depolarization versus Ba^2+^ current at the same time point.

We next utilized a FRET 2-hybrid binding assay (45,46) to evaluate the relative strength of interactions between the IQ and ICDI regions. FRET binding pairs were constructed by tagging cerulean fluorescent protein to ICDI_1.4_, and venus fluorescent protein to PreIQ_3_-IQ-A_1.3_, a peptide that includes IQ_1.3_ as well as ~30 residues upstream (PreIQ_3_) and ~150 residues downstream (A-region) of the IQ domain (**Fig. 1D, Fig. S1**). Both PreIQ_3_ and A regions were included initially to ensure that all likely interacting residues were included. Strong binding was detected between the venus-PreIQ_3_-IQ-A_1.3_ and Cerulean-ICDI_1.4_, as can be appreciated by the steep FRET binding curve determined by the FRET Ratio (FR) of each cell plotted as a function of the free donor concentration (Cerulean tagged ICDI_1.4_) (**Fig. 1D**, black). After calibration, the FRET binding curve for WT venus-PreIQ_3_-IQ-A_1.3_ *versus* Cerulean-ICDI_1.4_ yielded a *K*_a_ of 21.4 μM^-1^. To identify key residues that support a high-affinity IQ/ICDI interaction, we undertook systematic alanine substitution of the IQ domain and evaluated the effect on binding affinity in our FRET assay. Within IQ_1.3_, we substituted each residue with an alanine or, at loci where the wild-type channel featured an alanine, we replaced the residue with a threonine. For identification of each residue, the canonical isoleucine is assigned position 0. Application of our FRET assay to each mutated peptide identified three residues, Y[-5]A, F[-2]A and F[+4]A, which severely perturbed the IQ/ICDI interaction (**Fig. 1E, Fig. S1**). Focusing on F[-2]A, FRET binding produced a shallower curve as compared to WT (**Fig. 1D** blue vs. gray), resulting in a *K*_a_ of 5.8 μM^-1^ (**Fig. 1E**, blue). Introducing this mutation into the chimeric channel resulted in a partial rescue of CDI (**Fig. 1F**), indicating that this interaction site is functionally relevant. However, the IQ domain substitutions Y[-5]A and F[+4]A, which also had a marked effect on *K*_a_, resulted in minimal CDI rescue (**Fig. 1F, G, Fig. S2**). Importantly, these residues also serve as anchors for apoCaM binding to the Ca_V_1.3 IQ domain, resulting in weak baseline CDI even in the absence of the ICDI domain (**Table S1**) (13).

In order to confirm the functional relevance of each binding loci identified, we turned to a previously described analysis known as individually transformed Langmuir (iTL) analysis (14). This approach allows us to rigorously correlate relative changes in binding with functional changes in CDI. As iTL was initially derived to evaluate the binding interfaces critical to CaM mediated channel regulation (13,14), we adjust the model to reflect the competitive binding scheme between CaM and ICDI (20). To do so, we first account for the ambiguity caused by binding sites which are important for both apoCaM and ICDI binding. We therefore incorporate both the IQ domain’s intrinsic affinity for apoCaM (*K*_a-CaM_) and that for the ICDI segment (*K*_a-ICDI_), the competitive inhibitor, by adjusting our measured *K*_a-ICDI_ such that:

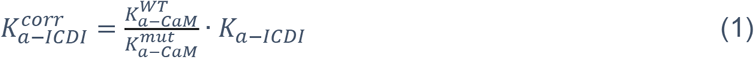

Where 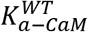 is the apoCaM binding affinity of WT PreIQ_3_-IQ-A_1.3_, and 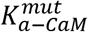 represents the apoCaM binding affinity of each mutant peptide, values which were previously measured (13) and are listed in **Table S1**. This compensation remains valid provided that the local concentration of ICDI is much greater than *K*_d-ICDI_ ([ICDI]>> 1/*K*_a-ICDI_). With this adjustment made, CDI can be defined by a modified Langmuir function as follows:

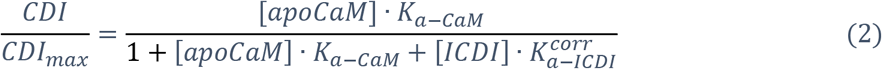

where *CDI* is the strength of CDI under endogenous levels of CaM; *CD/*_max_ is the CDI in saturating concentrations of CaM (**Table S1**); *K*_a-CaM_ is the association constant for apoCaM binding to PreIQ_3_-IQ-A_1.3_; [apoCaM] is the free apoCaM concentration in the cell; and [ICDI] is the effective local concentration of ICDI. Equation 2 predicts an inverse correlation between 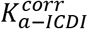 and relative *CDI*, which we can fit to our data (**Fig. 1H**). For channels containing WT IQ_1.3_ and ICDI_1.4_ domains, the strong binding between IQ and ICDI results in minimal CDI, as seen by the black data point on the plot. In contrast, Ca_V_1.3_s_ channels, which lack an ICDI domain such that 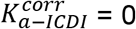 by definition, display large CDI values (cyan). The F[-2]A mutation, which produced a partial restoration of CDI, resides in an intermediate position (blue). Overall, our data can be well fit by equation 2 (**Fig. 1H**), confirming the functional relevance of the identified residues in a competitive model.

### Alanine scanning of the ICDI domain reveals complementary hotspots

Having identified several critical residues within the IQ domain required for ICDI binding, we next probed the ICDI for critical determinates of binding to the IQ. In order to scan a more extensive segment of the channel, we made triple alanine substitutions for every three contiguous residues within the ICDI domain. The ICDI domain has previously been localized to amino acids 1868-1956 of the Ca_V_1.3 DCT (29,32,33). We therefore undertook our alanine scan on this segment of the channel. Importantly, this region includes the distal C-terminal regulatory domain (DCRD), which was previously identified as playing an important role in the ICDI mediated inhibition of CDI (27,31,33,47). Finally, we included a selective v[1907]A mutation, as this amino acid change has previously been shown to dramatically alter the function of ICDI (30). To evaluate the effect of these alanine substitutions on IQ/ICDI binding, we again utilized our FRET two-hybrid binding assay, pairing venus-PreIQ_3_-IQ-A_1.3_ with Cerulean-ICDI_1.4_ (**Fig. 2A, Fig. S3**). Indeed, measured *K*_a_ values revealed multiple hotspots within the ICDI domain, with a wide range of binding affinities with the IQ containing peptide (**Fig. 2B**). The two mutation sites displayed in gray (KQE[1911]AAA and YSD[1941]AAA) were not evaluated as they failed to express. Notably, the effects of the hotspots on the ICDI domain were significantly larger than those observed within the IQ domain (**Fig. 2B** *vs.* **Fig. 1E**).

**Figure 2.**
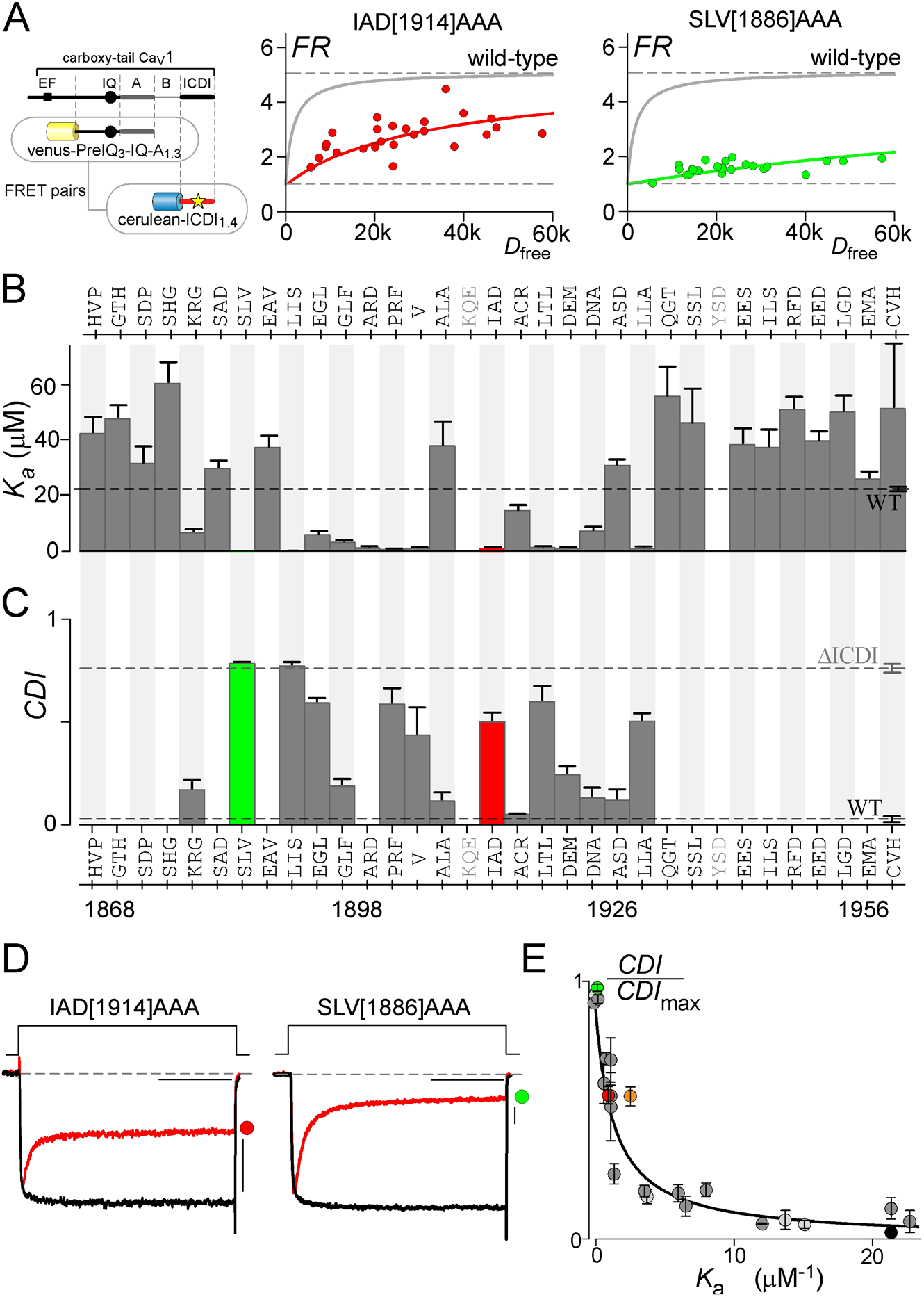
Identifying the hotspots on ICDI_1.4_ required for ICDI binding and function. (**A**) FRET binding partners are displayed on the left, with the star indicating the locus of alanine mutations within ICDI_1.4_. Compared to WT (gray line), IAD[1914]AAA (red) and SLV[1886]AAA (green) significantly decreased binding affinity between the FRET pairs to differing degrees. (**B**) Summary of *K*_a_ values for venus-PreIQ_3_-IQA_1.3_ *versus* mutant Cerulean-ICDI_1.4_ peptides measured with FRET two-hybrid as in panel A. Alanine was systematically substituted three amino acids at a time into Cerulean-ICDI_1.4_, with the identity of the amino acids displayed on the top of the bar graph. Amino acids labeled in gray (KQE, YSD) were not evaluated due to poor expression. The dashed line indicates the WT value and the colored bars correspond to data in other panels of the figure here and throughout. Data are displayed ± SEM. (**C**) Summary of CDI values measured for Ca_V_1.3_ΔDCT1.4_ harboring mutations corresponding to mutations which produced large effects on binding affinity in **B**. Gray dashed line represents the robust CDI of the channel without ICDI_1.4_, while the black dashed line indicates the nominal CDI of WT Ca_V_1.3_Δ/DCT1.4_. Data are displayed ± SEM. (n=3-8) (**D**) Exemplar whole cell recordings of Ca_V_1.3_Δ/DCT1.4_, demonstrating a significant restoration of CDI due to IAD[1914]AAA and near complete CDI due to SLV[1886]AAA. (**E**) Robust fit of CDI (**C**) and FRET binding data (**B**) to equation 2, using identical parameters as in Figure 1H. Light gray circles indicate data from Figure 1, while dark gray and colored circles indicate mutations within ICDI corresponding to **B** and **C**, with green and red circles indicating exemplars shown in other panels for reference. The PKA phosphorylation site Ser1883 is plotted as the orange circle.

In order to correlate loss of binding affinity with function, we measured the CDI those mutations which exhibited a large change in binding affinity (**Fig. 2C, Fig. S4**). As predicted, mutations which resulted in a significant loss of IQ/ICDI binding also exhibited a corresponding restoration of CDI. Focusing on two examples, IAD[1914]AAA moderately reduced IQ/ICDI binding (**Fig. 2A,B,** red), while introduction of the same mutations into our chimeric channel enabled a partial restoration of CDI (**Fig. 2C, D,** red). On the other hand, SLv[1886]AAA displayed a drastic reduction in IQ/ICDI binding (**Fig. 2A, B** green), and CDI was fully restored to the level seen in Ca_V_1.3_Δ1626_ (**Fig. 2C, D** green). Importantly, all identified loci are well fit by our Langmuir function, such that the same set of equation 2 parameters describes both the IQ region and ICDI (**Fig. 2E**). Of note, as mutations in ICDI do not affect the binding of apoCaM, the correction factor for *K*_a_ is no longer required, and 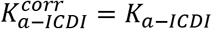. Having identified critical loci within ICDI we note that one of these amino acids (S[1866]) has previously been identified as a phosphorylation site which reduces the binding affinity of ICDI_1.4_ by about 10 fold, while increasing CDI of Ca_V_1.3_Δ1626/DCT1.4_ (43). We therefore included the results of phosphorylation at this site in our analysis (**Fig. 2E,** orange). Indeed, phosphorylation of this amino acid results in a change in binding affinity which correlates with CDI according to the same Langmuir function. Thus, we have identified numerous residues within ICDI_1.4_ which are critical determinants of a functional competition between ICDI and apoCaM for the IQ region of Ca_V_1.3.

### The role of the A region in the IQ/ICDI interaction

While our analysis identified several functionally-relevant IQ domain loci, the impact of these mutations was far less than those identified within the ICDI (**Fig. 1** *vs*. **Fig. 2**). This suggests that additional regions outside the IQ domain may contribute to ICDI binding. In order to identify such regions, we generated truncated variations of our venus*-*PreIQ_3_-IQ-A_1.3_ construct and paired them with Cerulean-ICDI_1.4_ in the FRET two-hybrid binding assays (**Fig. 3A, Fig. S5**). We began by removing the PreIQ_3_, and found no change in FRET binding, indicating that all relevant interaction loci are contained within the IQ and A regions (**Fig. 3B,** blue). However, removal of the IQ domain, leaving only the A region intact, resulted in a complete loss of FRET binding (**Fig. 3B,** open circles). Likewise, the IQ region alone displayed no binding with ICDI (**Fig. 3C, B)**, suggesting that both the IQ and the A region are necessary for interaction with ICDI. In order to further localize the critical interaction sites, we undertook successive truncation of the vernus-IQ-A_1.3_ peptide (**Fig. 3D**). FRET measurements demonstrated minimal effect of truncations up to 34 amino acids from the end of the A region, as demonstrated by the strong binding of venus-IQ-A_1.3_Δ34 with ICDI_1.4_ (**Fig. 3E, green**). However, our next truncation, venus-IQ-A_1.3_Δ28, exhibited a marked decrease in FRET binding (**Fig. 3E, F,** red) Thus, the 34 residues immediately downstream of the IQ domain critically augment ICDI binding. Of note, this region includes the previously identified proximal C-terminal regulatory domain (PCRD), which is reported to play an important role in the ICDI interaction (27,31,33,47). Having identified a subset of the A region which is vital to ICDI binding, we again undertook systematic alanine substitutions, replacing each 3 contiguous residues with three alanine residues and undertook our FRET-2-hybrid binding assay (**Fig. 3G**). Disruptions in binding were identified as the result of a number of mutations, spanning both the previously identified PCRD region as well as a previously unidentified region upstream of this motif (**Fig. 3H, I, Fig. S6**). Thus, both the IQ and the distal A region of the channel are required for high-affinity interaction with ICDI.

**Figure 3.**
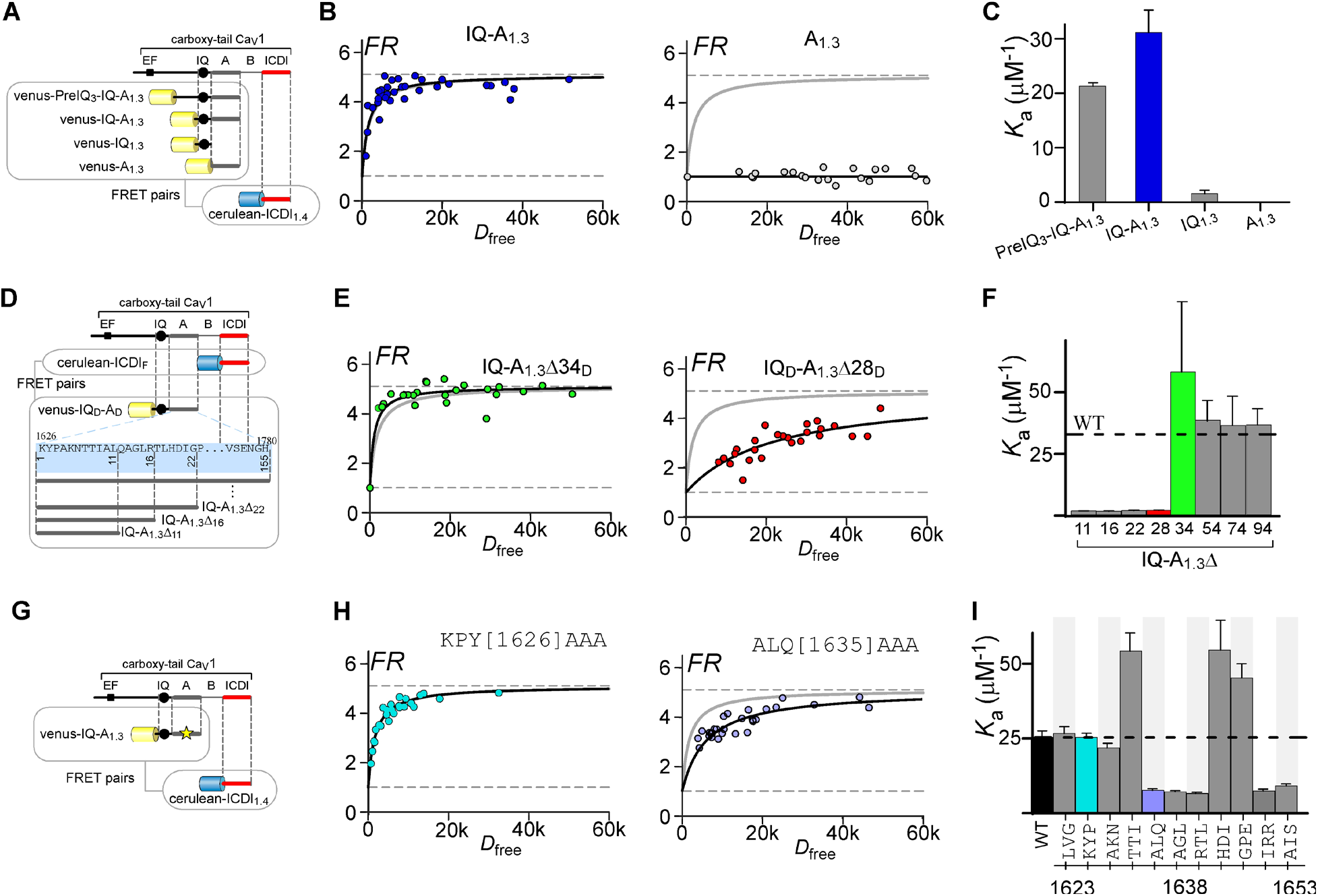
Elucidating the role of the A region for ICDI interaction. (**A**) Cartoon depicting FRET interacting pairs for panels **B** and **C**. Portions of the PreIQ_3_-IQ-A_1.3_ were paired with ICDI_1.4_ to identify critical regions. (**B**) IQ-A_1.3_ is sufficient to support robust FRET (blue), indicating that the pre-IQ3 region is not involved in the interaction. However, the A region alone is not sufficient to support binding (open circles). (**C**) *K*_a_ values for each channel fragment indicate that neither IQ_1.3_ or A_1.3_ is sufficient for robust binding with ICDI_1.4_, however both regions are necessary as indicated by the robust binding of IQ-A_1.3_ with ICDI_1.4_. Data are displayed as mean ± SEM here and throughout. (**D**) Cartoon depicting FRET interacting pairs for panels **E** and **F**. Cerulean-ICDI_1.4_ was paired with various truncations of Venus-PreIQ_3_-IQ-A_1.3_. the blue box shows the sequence of the A region and illustrates the truncation strategy. (**E, F**) Truncating up to 34 amino acids distal to the IQ region (Δ34) had nominal effects on binding (green), however removal of 6 additional amino acids (Δ28) dramatically reduced binding affinity (red). (**G**) Cartoon depicting FRET interacting pairs for panels **H** and **I**. Alanine substitutions were made within the A region, as indicated by the yellow star. (**H**) Mutation of KPY[1626]AAA within the A region had no effect on binding (cyan), while ALQ[1635]AAA (purple) diminished the binding affinity between IQ-A_1.3_ and ICDI_1.4_. (**I**) Summary of *K*_a_ values for alanine mutations in the A region indicate multiple critical amino acids.

### The functional relevance of ICDI in Ca_V_1.2 channels

Similar to Ca_V_1.3 and Ca_V_1.4, Ca_V_1.2 channels also feature a highly homologous ICDI segment, argued to function as a channel inhibitor (33,48) or as a transcriptional factor (41,42). We therefore considered the impact of ICDI_1.2_ on both Ca_V_1.2 and Ca_V_1.3 channels. We interrogated the binding of Cerulean-ICDI_1.2_ with venus-PreIQ_3_-IQ-A_1.2_ via FRET two-hybrid (**Fig. 4A, B**), and found that the interaction is significantly weaker than the prototypic venus-PreIQ_3_-IQ-A_1.2_ and Cerulean-ICDI_1.4_ interaction (**Fig. 4B** *vs*. **Fig. 1D**). However, when paired with venus-PreIQ_3_-IQ- A_1.3_, binding with ICDI_1.2_ is significantly larger, and only about half that of the strong binding of ICDI_1.4_ (**Fig. 4C**). Thus, it appears that ICDI_1.2_ is poised to have a larger effect in the context of Ca_V_1.3 channels as compared to its native channel backbone. Nonetheless, the limited binding observed between Cerulean-ICDI_1.2_ with venus-PreIQ_3_-IQ-A_1.2_ prompted us to evaluate the possibility of a functional role for ICDI_1.2_ within Ca_V_1.2 channels. Interestingly, truncation of Ca_V_1.2 at the known carboxy-tail cleavage site (49) for this channel (Ca_V_1.2_Δ11800_) resulted in a minimal, yet statistically significant (p ≤ 0.05), increase in CDI (**Fig. 4D, F, Fig. S7**). Next, to test the effect of ICDI_1.2_ on Ca_V_1.3 channels we replaced the native ICDI_1.3_ of Ca_V_1.3 long channels with ICDI_1.2_. Indeed, the loss of CDI surpassed that of Ca_V_1.2 channels (**Fig. 4E**), as predicted based on the stronger PreIQ_3_-IQ-A_1.3_ /ICDI_1.2_ interaction (**Fig. 4C**). In fact, the CDI exhibited by Ca_V_1.3-ICDI_1.2_ channels was not statistically different than the CDI measured in the native Ca_V_1.3_long_ splice variant (**Fig. 4F**).

**Figure 4.**
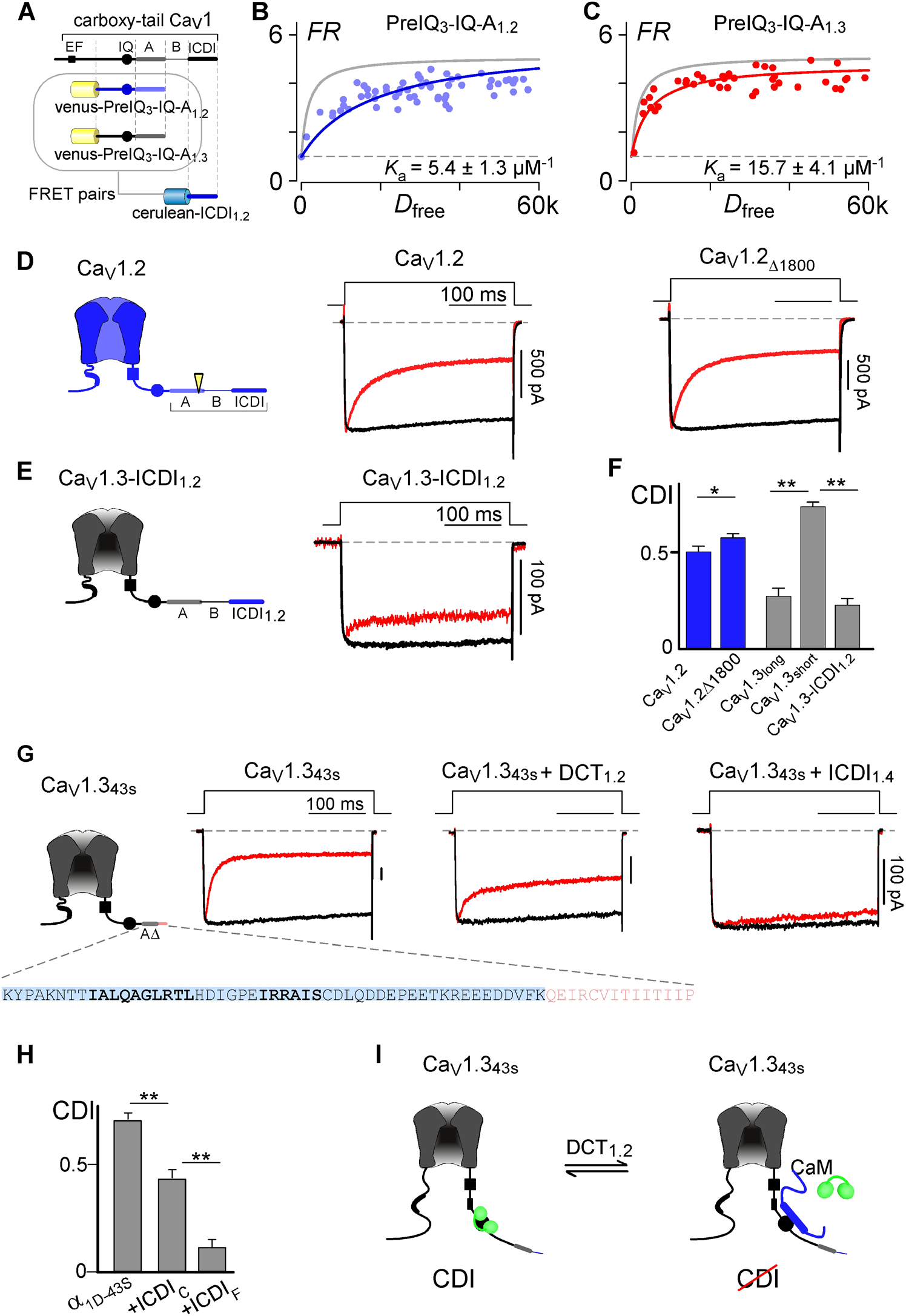
Residual functionality of ICDI1.2 in modulating Ca_V_ channels. (**A**) Cartoon depicting FRET interacting pairs for panels B and C. Venus-PreIQ_3_-IQ-A_1.2_ or Venus- PreIQ_3_-IQ-A_1.3_ was paired with Cerulean-ICDI_1.2_ in order to evaluate the role of the ICDI contained within Ca_V_1.2 channels. (**B**) Venus-PreIQ_3_-IQ-A_1.2_ displays moderate binding with Cerulean- ICDI_1.2_ (blue) as compared to the robust binding between Venus-PreIQ_3_-IQ-A_1.3_ and Cerulean- ICDI_1.4_ which is reproduced as the gray line for comparison. (**C**) Venus-PreIQ_3_-IQ-A_1.3_ is able to bind strongly with Cerulean-ICDI_1.2_. (**D**) *Left:* channel cartoon with cleavage site indicated by the yellow arrowhead. *Right:* Exemplar data demonstrating that truncation of Ca_V_1.2 at the cleavage site has a minor effect on CDI. (**E**) *Left:* channel cartoon indicating introduction of ICDI_1.2_ into the Ca_V_1.3 channel backbone. *Right:* Introduction of ICDI_1.2_ into Ca_V_1.3 channels causes a large decrease in CDI. (**F**) Average CDI data demonstrating a modest but statistically significant effect of ICDI_1.2_ on Ca_V_1.2 channels (blue), and a large effect on Ca_V_1.3 which is comparable to the native effect of ICDI1.3 contained within Ca_V_1.3_long_ (gray). Data are plotted as mean ± SEM. (*p≤0.05, ** p≤0.01) (**G**) *Left:* cartoon of human Ca_V_1.3_43S_, with the sequence of the end of the channel displayed below. This splice variant contains a portion of the A region (blue highlight) which contains the identified hotspots required for ICDI binding (bold), as well as a short sequence of unique amino acids prior to truncation of the channel (pink). *Right:* exemplar whole cell patch clamp data demonstrates robust CDI in WT Ca_V_1.3_43S_, which is reduced when the predicted cleavage fragment of the human Ca_V_1.2 channel is co-expressed, or when ICDI_1.4_ is expressed as a peptide. (**H**) Average CDI effects for the channels described in panel G, DCT_1.2_ and ICDI_1.4_ both have significant effects on the CDI of Ca_V_1.3_43S_ (** p≤0.01, data are displayed ± SEM). (**I**) Cartoon proposing a cross-channel effect of ICDI_1.2_, such that cleaved or independently transcribed DCT_1.2_ may modulate Ca_V_1.3_43S_ channels.

Multiple studies have shown that the DCT of Ca_V_1.2, containing ICDI_1.2_, exists as a peptide within neurons and cardiomyocytes, either due to proteolysis (33,49), or as a result of alternative transcriptional initiation sites (41). Moreover, it has previously been demonstrated that ICDI domains can exert their effects on L-type channels when expressed as separate peptides (27). We therefore sought to recreate the potential interaction of select Ca_V_1.3 channel variants with the DCT of Ca_V_1.2. To begin, we choose the human Ca_V_1.3_43S_ splice variant of Ca_V_1.3, as these channels terminate just past the A region and thus lack an inherent ICDI module (27). In addition, the inclusion of the A region within Ca_V_1.3_43S_ has been demonstrated to be important for ICDI binding, both in functional experiments done by others (27) and in our alanine scan of the A region (**Fig. 3**). We therefore generated the proteolytic product of human Ca_V_1.2 channels (DCT_1.2_), and evaluated the effect of this peptide on the CDI of Ca_V_1.3_43S_. Indeed, consistent with previous studies (27), co-expression of DCT_1.2_ significantly reduced the CDI of Ca_V_1.3_43S_ (**Fig. 4G, Fig. S7**). For comparison, we also co-expressed these channels with ICDI_1.4_ expressed as a peptide, which we have shown has a *K*_a_ about double that of ICDI_1.2_ (**Fig. 4C** *vs*. **Fig. 1D**). Indeed, ICDI_1.4_ results in an even larger CDI deficit when expressed with Ca_V_1.3_43S_ (**Fig. 4G**). Thus, the proteolytically cleaved DCT_1.2_ is well poised to exert a significant modulation of select Ca_V_1.3 channel variants, such that the ambient concentration of CaM and DCT_1.2_ are able to tune the CDI of Ca_V_1.3_43S_ channels in a competitive manner (**Fig. 4H**).

## Discussion

CaM regulation of Ca_V_ channels is vital to normal physiology, and thus has been the subject of intense study (8,10,11,17,50–52). The competitive mechanism of ICDI within select Ca_V_1 channels forms a basis with which CaM regulation can be tuned (12,20,32). Numerous processes designed to modulate this regulation include splice variation, RNA editing, variations in ambient CaM concentration and phosphorylation (12,13,18,36,43). Identification of critical loci involved in this regulation is therefore key to understanding how CaM regulation may vary in different physiological and pathological states. As such, in-depth residue-level analysis not only reveals interfaces utilized by cells to tune channel regulation, but may offer targets in the search for novel regulators of the channel which may have therapeutic benefit. In particular, the dramatically different efficacy of ICDI across channel subtypes may offer the possibility of subtype selective drug targeting, which remains challenging for Ca_V_1 channels.

Given the importance of understanding these interactions within the C-tail of Ca_V_1 channels, we quantified the structure-function relationship of these interactions using a variant of previously described iTL analysis (14). This provided a major advantage in that the quantitative agreement of our results with equation 2 demonstrates that each identified locus is functionally relevant. This overcomes a common limitation of binding assays between channel fragments, which may identify sites which are inaccessible or inconsequential in the context of the holochannel. Moreover, by fitting to a specific Langmuir curve, we can distinguish mutations which may alter channel function through ancillary mechanisms such as transduction or altered folding of the channel. Thus, in addition to identifying critical loci, our results confirm the competitive mechanism described for ICDI modulation of CDI.

A number of previous studies have identified regions within the C-tail of Ca_V_1 channels which are critical to the competitive mechanism of ICDI inhibition. Among these are the PCRD and DCRD regions, which were identified within Ca_V_1.2 channels as potential interaction sites such that the DCRD region of the proteolytically cleaved Ca_V_1.2 DCT may interact with the PCRD on the channel via electrostatic interaction with the negatively charged amino acids (33). Homologous regions within Ca_V_1.3 and Ca_V_1.4 were later shown to be important for ICDI regulation, and for the interaction with modular peptides (27,30,31,33,47). In this study, the PCRD resides within the A region, and overlaps with the identified locus of critical amino acids required for high affinity binding between the IQ-A and ICDI. Interestingly, these critical amino acids were identified both within the PCRD, as well as upstream of the motif, arguing for a larger interacting region which is highly conserved across Ca_V_ channels (**Fig. S8**). However, our binding assay also demonstrated that the A region, in itself, is insufficient for high affinity binding, but also requires the upstream IQ region (**Fig. 3A**). This fits with previous findings in which neutralization of the PCRD arginines was not sufficient to prevent the ICDI inhibition of CDI (31), pointing to the existence of additional interaction loci. In a similar manner, our scan of the ICDI region validated the importance of the DCRD, while also identifying numerous critical interacting loci upstream of the motif (**Fig. S8**). Thus, this study has expanded our knowledge of the important interactions required for ICDI inhibition, providing a comprehensive map of the critical loci within the C-tail. The impact of ICDI in Ca_V_1.3 and Ca_V_1.4 channels has been well recognized, however its role in Ca_V_1.2 has been uncertain. It had been demonstrated that the truncation of Ca_V_1.2 results in increased current density, altered voltage dependence of channel activation and disrupted targeting of the channel to the membrane (33,53,54), however no impact on CDI has been reported. Here, we find that the impact of ICDI_1.2_ within Ca_V_1.2 channels is minimal, allowing cleavage of this DCT region without significant disruption of CDI. Yet ICDI_1.2_ is capable of causing significant disruption of CDI in the context of Ca_V_1.3 (**Fig. 4E-I**) (27). This selective effect of ICDI_1.2_ on Ca_V_1.3 channels is intriguing as it represents a difficult to achieve discrimination between Ca_V_1.2 and Ca_V_1.3 (55,56). The strong homology between these two channels and has resulted in challenges to dissecting the contribution of each channel to the function of cells which express both channel subtypes, and hinders therapeutic options for neuropsychiatric disorders which may benefit from blockade of Ca_V_1.3 (55). As such, the interface defined in this study between the IQ-A region and ICDI may represent a promising interface with which to selectively target Ca_V_1.3.

The DCT of Ca_V_1.2 is routinely cleaved in neurons and myocytes, with the cleavage product able to either remain associated with the channel (33), or translocate to the nucleus (41). Our FRET binding data would argue that this association between Ca_V_1.2 and the DCT may be relatively weak, leaving DCT_1.2_ available to other binding partners. Moreover, it has been shown that an alternate start site exists within the C-tail of Ca_V_1.2, such that alternative transcription of Ca_V_1.2 will produce a DCT peptide containing a calcium channel associated transcription regulator (CCAT) (41). Importantly, ICDI would be intact within this peptide, providing an additional source of DCT_1.2_ within cells. Our FRET binding analysis (**Fig. 4A**) suggests that the Ca_V_1.2 DCT is capable of binding upstream calmodulatory elements in Ca_V_1.2, albeit weakly. Functional analysis, however, suggests only minimal effects of this segment on Ca_V_1.2 CDI. By comparison, the Ca_V_1.3_43S_ channel variant contains all the elements required for high affinity binding with DCT_1.2_ and exhibits functional inhibition of CDI (**Fig. 4G**), consistent with previous studies (27). Thus, it seems likely that DCT_1.2_ may interact with this channel, altering the normally robust CDI. It is interesting to note, however, that while DCT_1.2_ is poised to modulate some Ca_V_1.3 channels, the same cannot be said of DCT1.3. Not only is the IQ-A region of Ca_V_1.2 suboptimal for binding to ICDI, but there is little evidence that Ca_V_1.3 is cleaved in neurons (31). Thus, this cross-channel modulation may be unidirectional. Finally, since Ca_V_1.2 and Ca_V_1.3 often exist within the same neuron, this mode of cross-channel modulation may represent an important method for tuning CDI in different regions of the brain.

### Experimental procedures

#### Molecular biology

The rat brain Ca_V_1.3 α1 subunit (in pcDNA6) corresponds to AF370009.1 (57), and was incorporated to the mammalian expression plasmid pCDNA6 (Invitrogen) (20). This plasmid features a unique *Bgl*II restriction site at a locus corresponding to ~450 amino acids upstream of the IQ domain, and a unique *XbaI* site after the stop codon which were used for generation of mutant and chimeric plasmids as descried below. The Ca_V_1.2 α1 subunit (in pGW) is identical to rabbit NM001136522 (58), and the Ca_V_1.4 channel (in pcDNA3) is the human clone corresponding to NP005174.2. Ca_V_1.3_Δ/DCT1.4_ was made by fusing with the DCT of the Ca_V_1.4 α1 subunit to the Ca_V_1.3 α1 subunit (truncated after the IQ domain), as previously described (20). Ca_V_1.3_43S_ was made by PCR amplification of the channel segment between the *BglII* site and IQ domain with the appendage of amino acids unique to this splice variant (27). The PCR product was then inserted into the channel via the *BglII/XbaI* sites.

FRET constructs were fluorescent-tagged (either Venus or Cerulean) using similar strategies as previously described (45). Briefly, Venus and Cerulean fluorophores (a kind gift from Dr. Steven Vogel, NIH) were subcloned into the pcDNA3 vector via unique *KpnI* and *NotI* sites. The PCR-amplified channel peptides, as described in Liu *et. al,* were then cloned in via unique *Not*I and *Xba*I sites (20). Mutations were introduced into the channel or FRET construct via PCR amplification or overlap extension PCR.

#### Transfection of HEK293 cells

For electrophysiology experiments, HEK293 cells were cultured on 10-cm plates, and channels were transiently transfected by a calcium phosphate protocol (10). We applied 8 μg of plasmid DNA encoding the desired pore forming α1 subunit, as well as 8 μg of β_2A_ (M80545) and 8 μg of rat α_2δ_ (NM012919.2) subunits along with 3 μg of SV40 T antigen. For microscope-based FRET assays, HEK293 cells cultured on 3.5-cm culture dishes with integral No. 0 glass coverslip bottoms (In Vitro Scientific) were transiently transfected using polyethylenimine (PEI) reagent (Polysciences).

#### Whole-cell patch clamp recordings

Whole-cell recordings were obtained using an Axopatch 200A amplifier (Axon Instruments). Electrodes were pulled from borosilicate glass capillaries (Word Precision Instruments), with 1-3 MΩ resistances, which were in turn compensated for series resistance by >60%. Currents were low-pass filtered at 2 kHz before digital acquisition at five times the frequency. A P/8 leak subtraction protocol was used. The internal solution contained (in mM): CsMeSO3, 114; CsCl, 5; MgATP, 4; HEPES (pH 7.4), 10; and BAPTA (1,2-bis(*o*- aminophenoxy)ethane- *N,N,N’,N’*-tetraacetic acid), 10; at 295 mOsm adjusted with CsMeSO3. The bath solution contained (in mM): TEA-MeSO3, 102; HEPES (pH 7.4), 10; CaCl2 or BaCl2, 40; at 305 mOsm adjusted with TEA-MeSO3. Data was analyzed using custom Matlab scripts. Inactivation was quantified as the ratio of current remaining after 300 ms in either Ca^2+^ or Ba^2+^ (r_300_), allowing CDI to be quantified as the difference between the r_300_ in Ba^2+^ vs Ca^2+^, measured at 10 mV for Ca_V_1.3 channels, and 30 mV cof Ca_V_1.2.

#### FRET optical imaging

FRET two-hybrid experiments were performed on an inverted microscope as described (45,46). The bath solution was a Tyrode’s solution composed of (in mM): NaCl, 138; KCl, 4; MgCl2, 1; HEPES (pH 7.4), 10; CaCl2, 2; at 305 mOsm adjusted with glucose. Background fluorescent signals were measured from cells without expression of the fluorophores and subtracted from cells expressing the fluorophores. Concentration-dependent spurious FRET was subtracted from the raw data prior to binding-curve analysis (45,46). Cerulean (59) and Venus (60) were used as the donor and acceptor fluorescent proteins instead of eCFP and eYFP, as their optical properties provided more robust and stable FRET signals. Acceptor-centric measurements of FRET were obtained with the 3^3^-FRET algorithm (45,46), in which the effective FRET efficiency (*E*_EFF_) and FRET ratio (*FR*) are defined as:

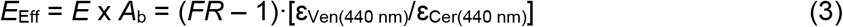

where *E* is the FRET efficiency of a donor-acceptor pair, *A*_b_ is the fraction of acceptor molecules bound by a donor, and ε_Ven(440 nm)_/ε_Cer(440 nm)_ is the approximate molar extinction coefficients of Cerulean and venus, which was measured as 0.08 on our setup. Intensity measurements at each wavelength were measured from individual cells such that variable expression across the cells enabled population of a binding curve. Binding curves were analyzed using GraphPad software (Prism), providing relative *K*_d_ values and standard error based on an unbiased the fit to the data. These relative Kd values were then calibrated according to a previously determined calibration factor (13,20), and converted to *K*_a_ = 1/*K*_d_. Importantly, the previous calibration factor determined for our setup utilized CFP/YFP FRET pairs. In order to account for the difference in FR using cerulean and venus, we determined the relative *K*_d_ for multiple peptides using both the florouphore pairs and found that the two data sets differed by a factor of 1.8, which we incorporated into the calibration factor.

## Data availability

All data is contained within the manuscript.

## Acknowledgements

This project was initiated under the direction of Dr. David Yue, who passed away in 2014. David was a brilliant scientist and exceptional mentor, and we are grateful for his guidance and friendship. We would like to thank Dr. Gordon Tomaselli for his guidance and support. We would also like to thank Hojjat Bazzazi for providing FRET plasmids, and for useful discussion and advice throughout the project. In addition, we thank Wanjun Yang for dedicated technical support and thank members of the Calcium Signals lab at Johns Hopkins for discussion and support of this project, as well as members of Dr. Dick’s lab at the University of Maryland for insightful discussions and editing of the manuscript.

## Funding and additional information

This grant was supported by an NIH/NINDS grant 5R01NS085074 and by an NIH/NHLBI grant 1R01HL149926.

## Conflict of interest

The authors declare that they have no conflicts of interest with the contents of this article.

## Abbreviations and nomenclature

CaM: calmodulin
apoCaM: Ca^2+^ free calmodulin
CDI: calcium/CaM dependent inactivation
Ca_V_: voltage gated calcium channel
Ca^2+^: calcium
FRET: fluorescence resonance energy transfer
FR: fret ratio
iTL: individually transformed Langmuir
CFP: cerulean fluorescent protein
YFP: yellow fluorescent protein
DCT: distal carboxy tail
ICDI: inhibitor of CDI
PCRD: proximal C-terminal regulatory domain
DCRD: distal C-terminal regulatory domain

## Notes

### Competing Interest Statement

The authors have declared no competing interest.

