## Supplementary Material for "Resolving the molecular fingerprint of the distal carboxy tail in modulating Ca_V_1 calcium dependent inactivation"

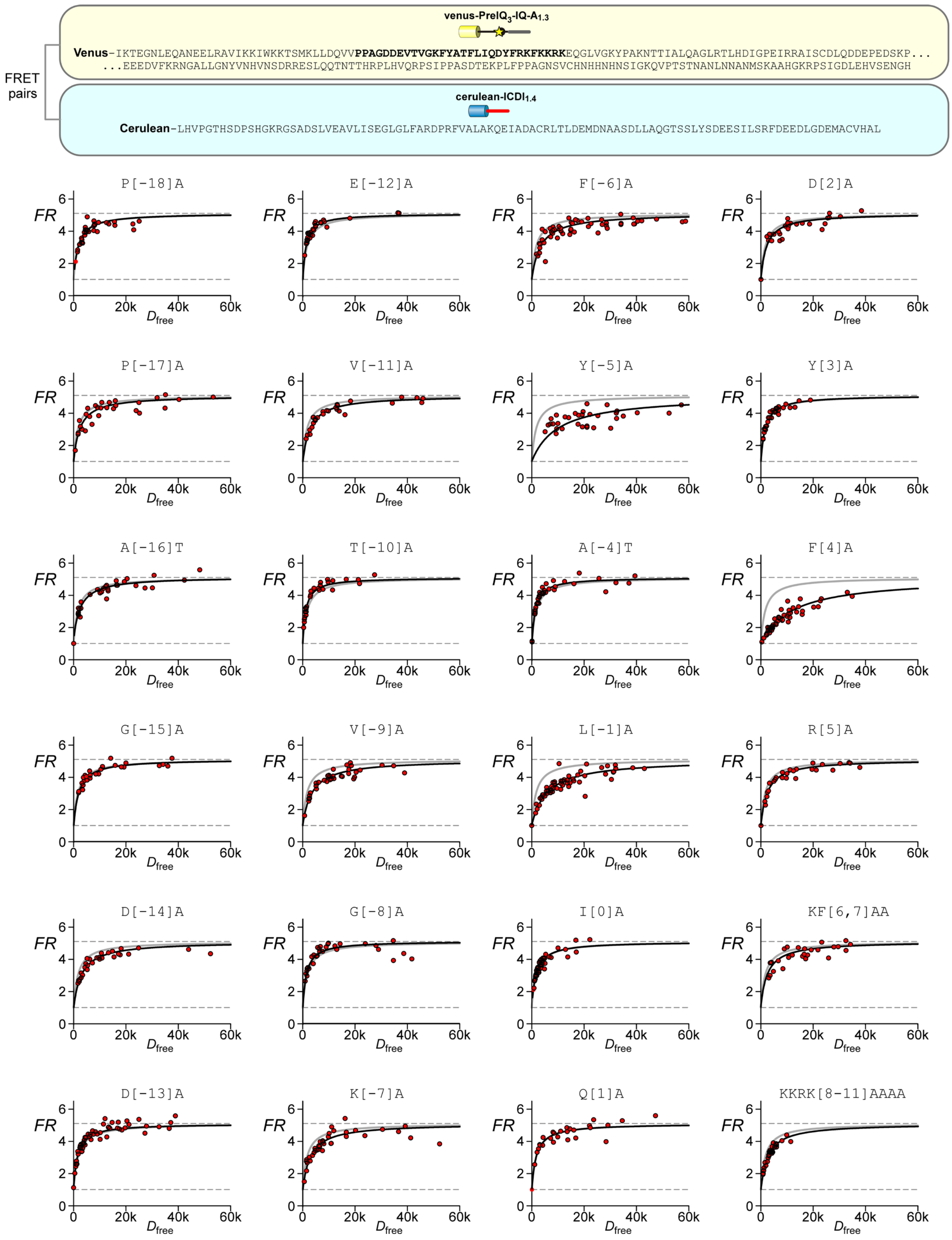


**Figure S1:** Top**:** Full sequence for the FRET 2-hybrid peptides used in main text Figure 1, with IQ region in bold. Below: FRET 2-hybrid binding curves for each mutation summarized in main text Figure 1E.

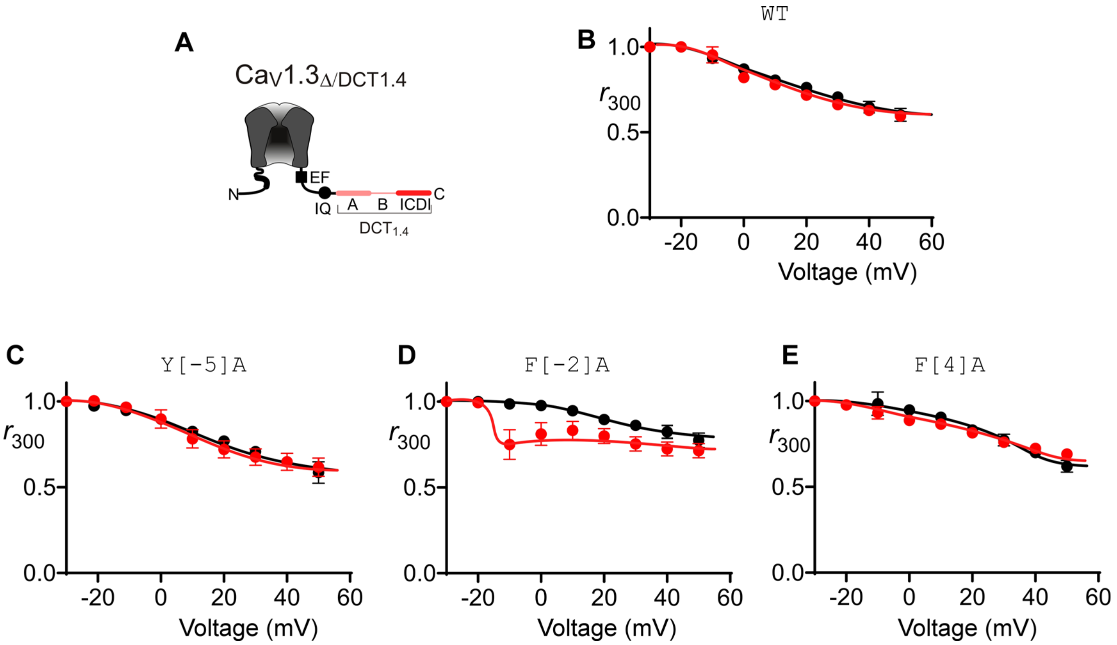


**Figure S2:** Full CDI dataset showing inactivation in $Ca^{2+}$ (red) versus $Ba^{2+}$ (black) as a function of voltage for WT and mutant $Ca_V1._{3\Delta/DCT1.4}$, corresponding to data displayed in Figure 1G of the main text. Data is displayed ± SEM, with n = 4-6.

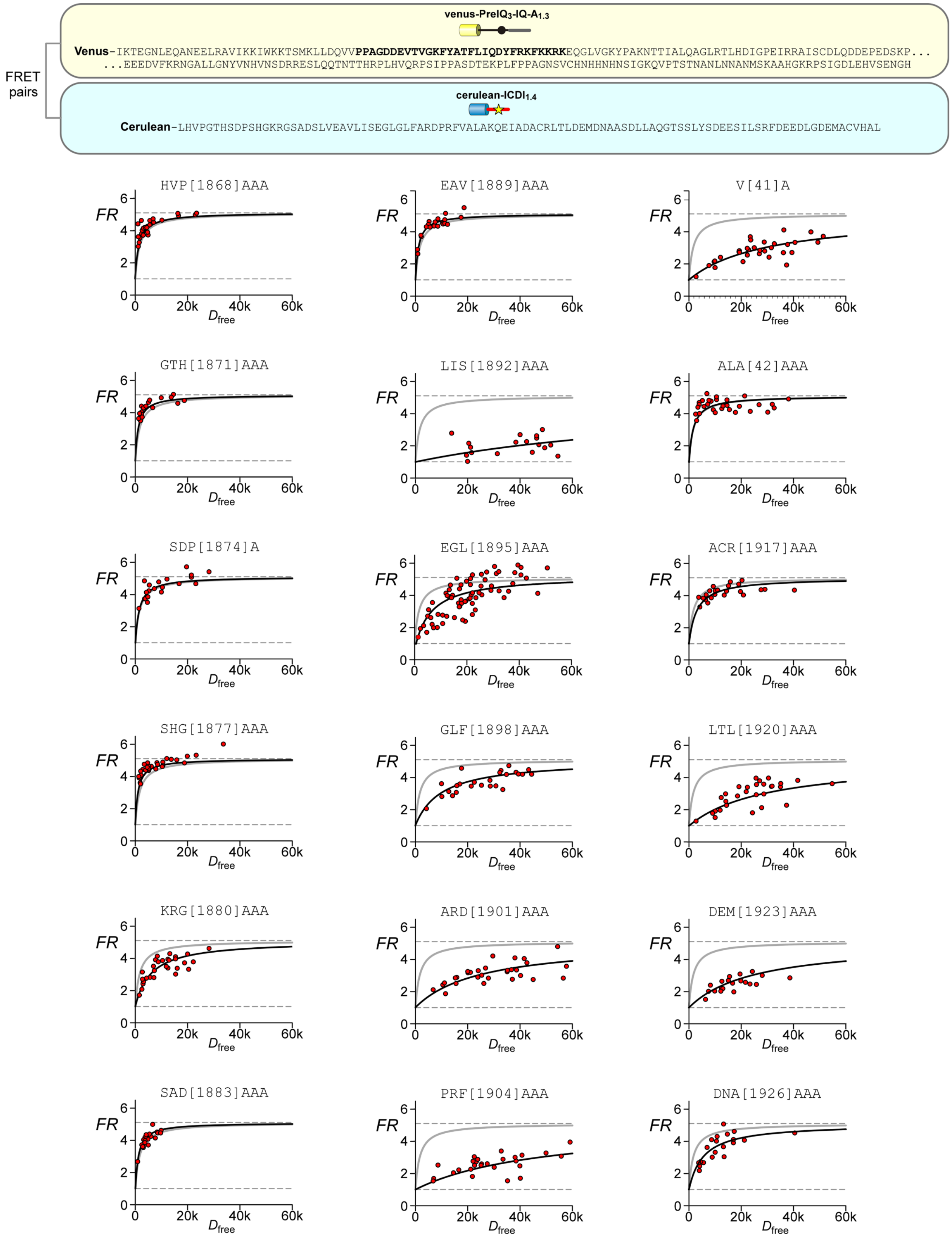


**Figure S3:** Top: Full sequence for the FRET 2-hybrid peptides used in main text Figure 2, with IQ region in bold. Below: FRET 2-hybrid binding curves for each mutation summarized in main text Figure 2B.

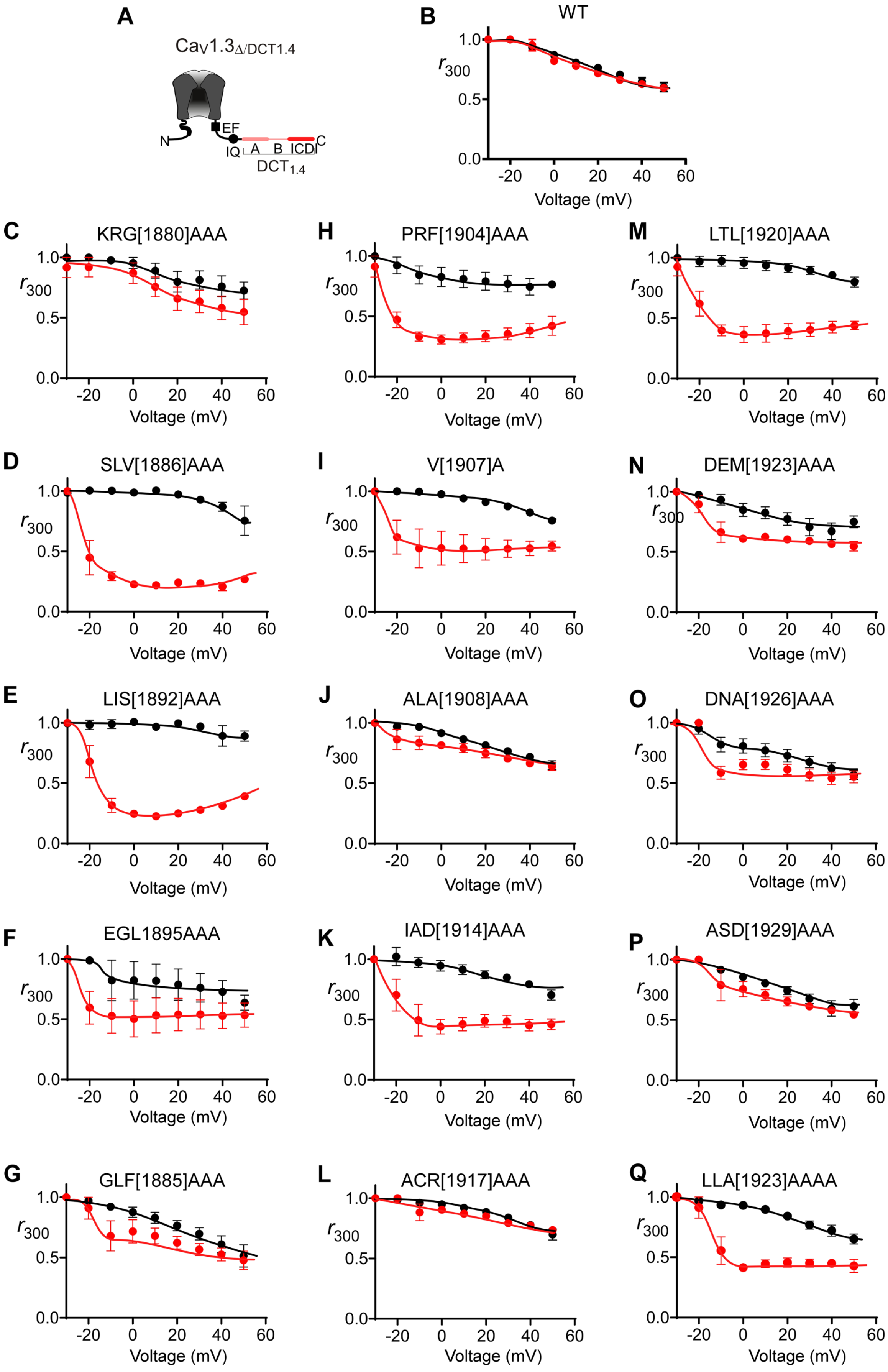


**Figure S4:** Full CDI dataset showing inactivation in $Ca^{2+}$ (red) versus $Ba^{2+}$ (black) as a function of voltage for WT and mutant $Ca_V1._{3\Delta/DCT1.4}$, corresponding to data displayed in Figure 2C of the main text. Data is displayed ± SEM, with n = 3-6.

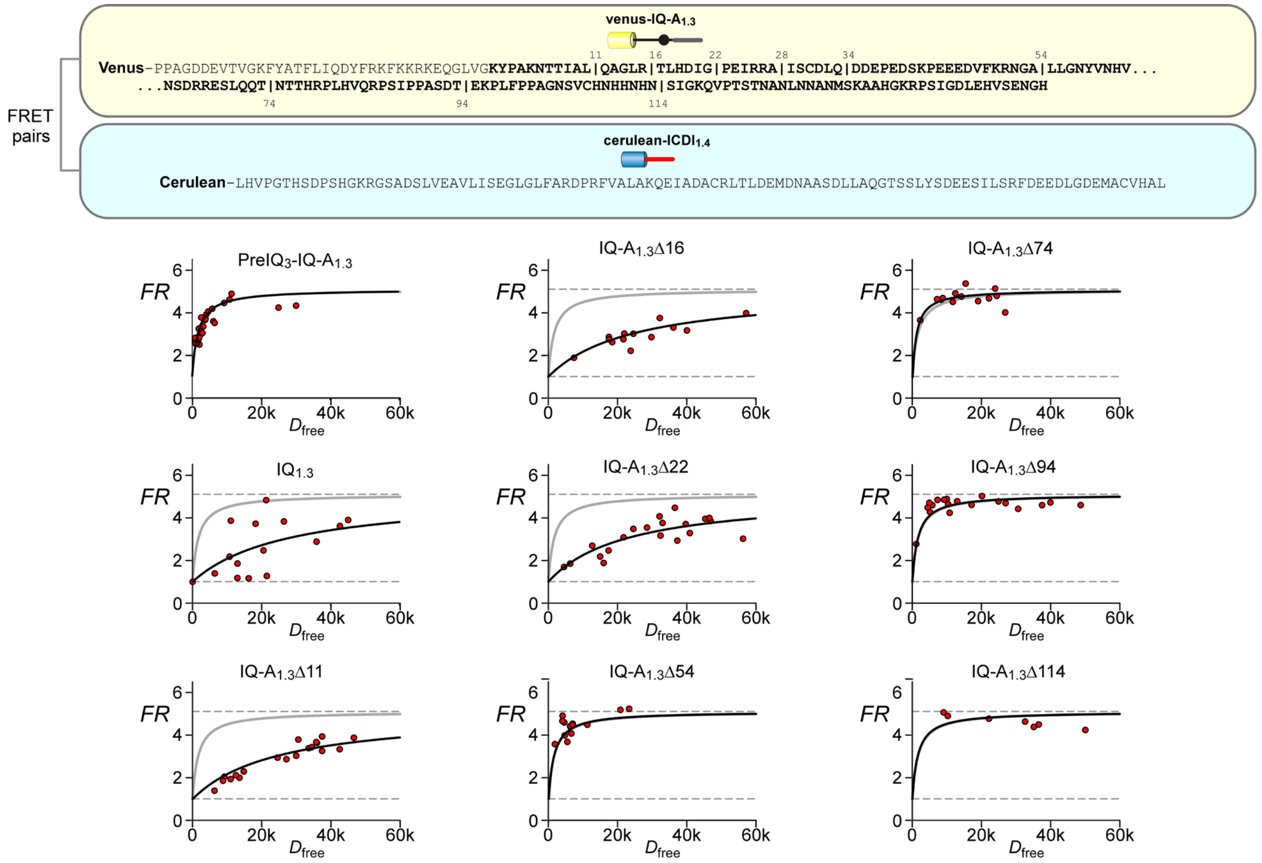


**Figure S5:** Top**:** Full sequence for the FRET 2-hybrid peptides used in main text Figure 3A-F, with A region in bold, and deletion sites marked by '|'.  Below: FRET 2-hybrid binding curves for each deletion summarized in main text Figure 2C,F.

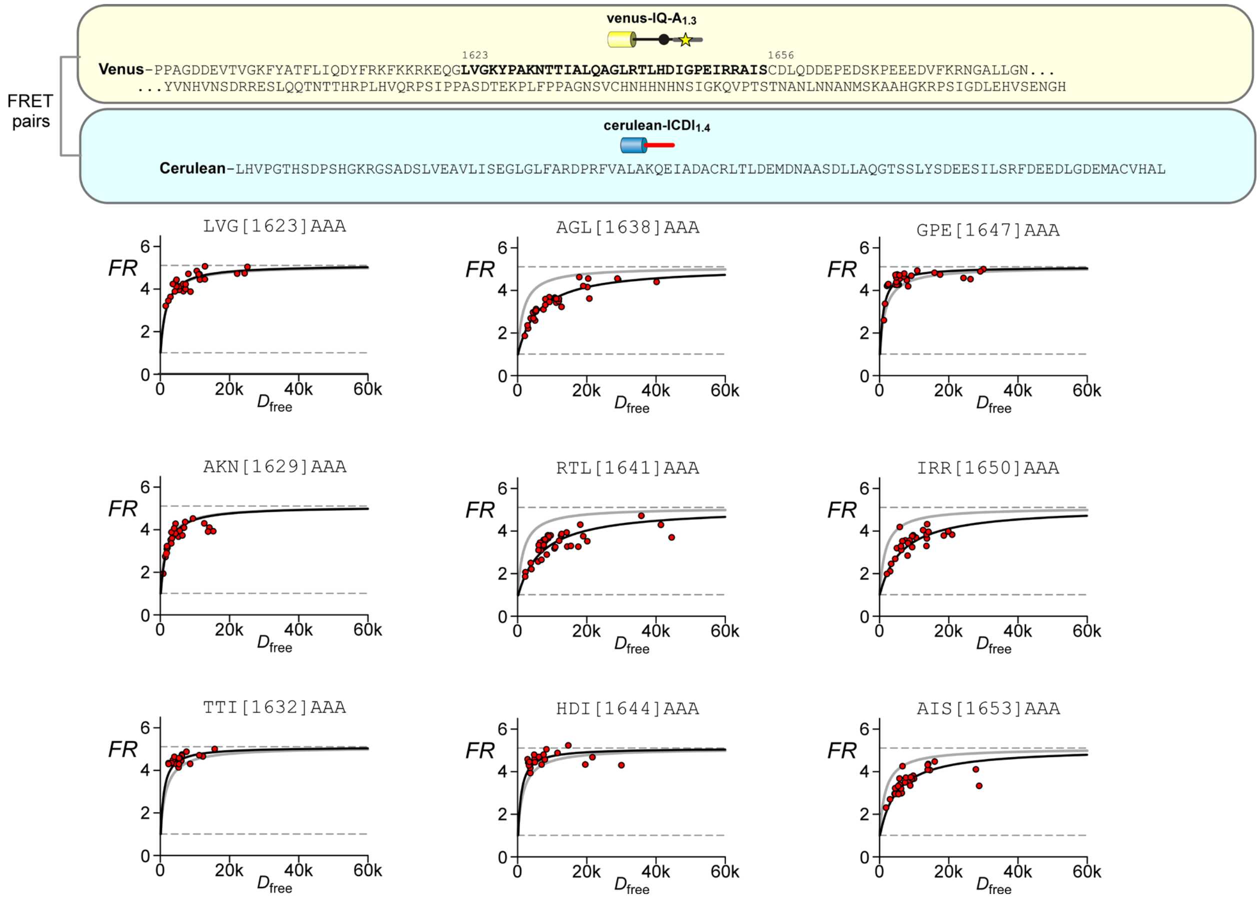

**Figure S6:** Top**:** Full sequence for the FRET 2-hybrid peptides used in main text Figure 3G-I, with the identified critical portion of the A region in bold. Below: FRET 2-hybrid binding curves for each mutation summarized in main text Figure 2I.

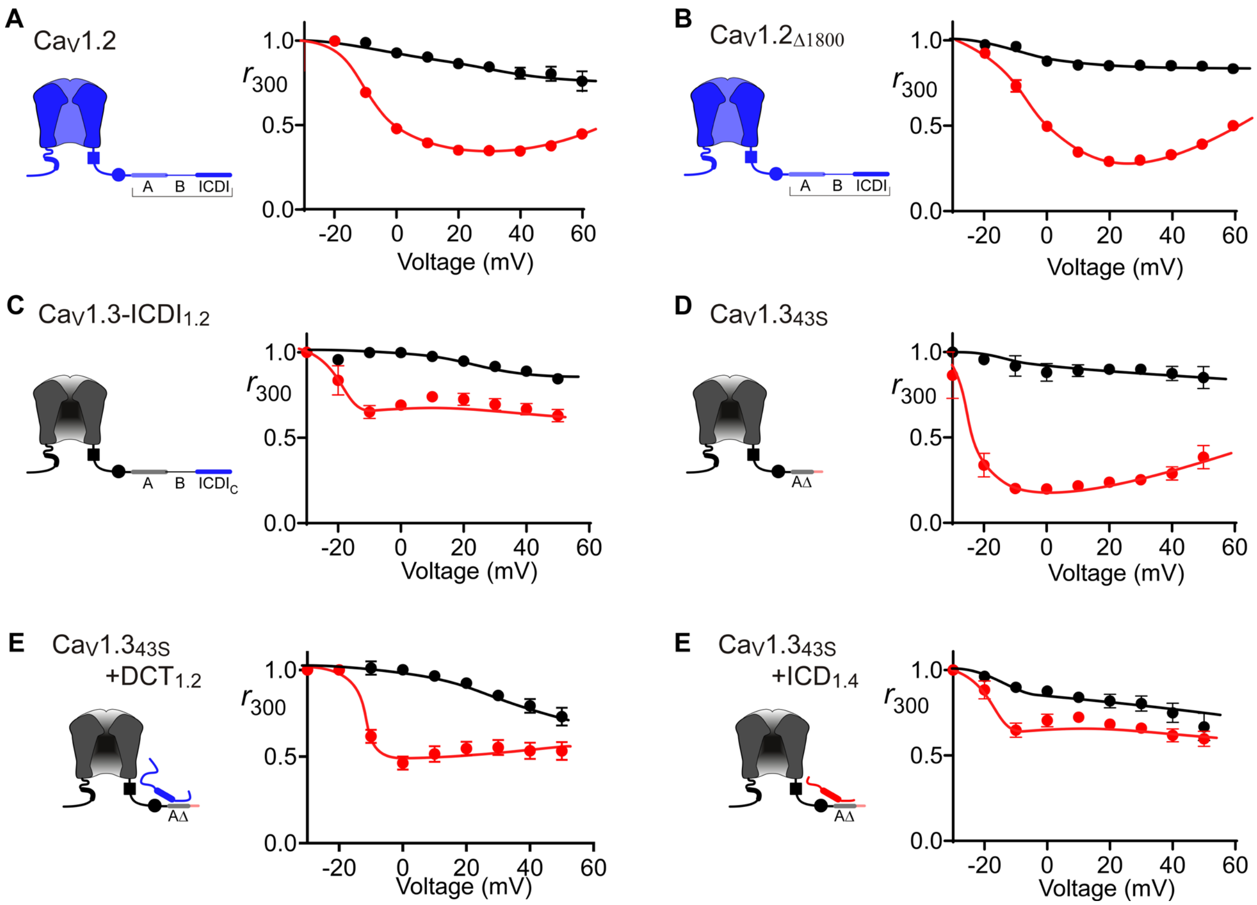


**Figure S7:** Full CDI dataset showing inactivation in $Ca^{2+}$ (red) versus $Ba^{2+}$ (black) as a function of voltage, corresponding to data displayed in Figure 4D-G of the main text. Data is displayed ± SEM, with n = 4-8.

```
                 preIQ3                                      IQ
CaV1.2 IKTEGNLEQANEELRAIIKKIWKRTSMKLLDQVVPPAGDDEVTVGKFYATFLIQEYFRKFKKRKEQGLV 1670
CaV1.3 IKTEGNLEQANEELRAVIKKIWKKTSMKLLDQVVPPAGDDEVTVGKFYATFLIQDYFRKFKKRKEQGLV 1624
CaV1.4 IKTEGNLEQANQELRIVIKKIWKRMKQKLLDEVIPPPDEEEVTVGKFYATFLIQDYFRKFRRRKEKGLL 1594

                                      A region
CaV1.2 GK-PSQRNALSLQAGLRTLHDIGPEIRRAISGDLTAEEELDKAMKEAVSAASEDDIFRRAGGLFGNHVS 1738
CaV1.3 GKYPAKNTTIALQAGLRTLHDIGPEIRRAISCDLQDDEPED-------SKPEEEDVFKRNGALLGNYVN 1686
CaV1.4 GNDAAPSTSSALQAGLRSLQDLGPEMRQALTCDTEEEEE------EGQEGVEEED-------------- 1643

                                      A region
CaV1.2 YYQSDSRSAFPQTFTTQRPLHISKAGNNQG---DTESPSHEKLVDS-TFTPSSYSSTG------SNANI 1797
CaV1.3 HVNSDRRESLQQTNTTHRPLHVQRPSIPPAS--DTEKPLFPPAGNSVCHNHHNHNSIGKQVPTSTNANL 1753
CaV1.4 ----------EKDLETNKATMVSQPSARRGSGISVSLPVGDRLPDSLSFGPSDDDRGT---PTSSQPSV 1699

                 A region                              B region
CaV1.2 NNANNTALGRLPRPAGYPSTVSTVEGHGSPLSPAVRAQEAAWKLSSKRCHSQESQIAMACQEGASQDDN 1866
CaV1.3 NNANMSKAAHGKRPSIGDLEHVSENGHYSYKHDRELQRRSSIKRTRYYETYIRSESGDEQLPTICREDP 1822
CaV1.4 PQAGSNTHRRGS---GALIFTIPEEGNSQPKG-------------------------TKGQNKQDEDEE 1740

                                      B region
CaV1.2 YDVRIGEDAECCSEPSLLSTEMLSYQDDENRQ------------------------------LAPPEEE 1906
CaV1.3 EIHGYFRDPRCFGEQEYFSSEECC-EDDSSPTWSRQNYSYYNRYPGSSMDFERPRGYHHPQGFLEDDDS 1890
CaV1.4 V-------------PDRLS-----YLDEQAGTPPCSV-------------------------LLPPHRA 1766

                                      B region
CaV1.2 KR--DIRLSPKKGFLRSASLG-RRASFHLECLKRQKNQGGD-----ISQKTVLPLHLVHHQALAVAGLS 1966
CaV1.3 PIGYDSRRSPRRRLLPPTPPSHRRSSFNFECLRRQNSQDDVLPSPALPHRAALPLHLMQQQIMAVAGLD 1959
CaV1.4 QRYMDGHLVPRRRLLPPTPAG-RKPSFTIQCLQRQGSCED------------LPIPGTYHR-------- 1814

                                      B region
CaV1.2 PLLQRSHSPTSLPRPCATPPATPGSRGWPPQPIPTLRLEGADSSEKLNSSFPSIHCGSWSGENSPCRGD 2035
CaV1.3 SSKAQKYSPSHSTRSWATPPATPPYRDWTPCYTPLIQVDRSESMDQVNGSLPSLHRSSWYTDEPDI--- 2025
CaV1.4 ---GRNSGPNRAQGSWATPPQ----RG-RLLYAPLLLVEEGAAGEYLGR-----------SSGP---- 1860

                                         ICDI
CaV1.2 SSAARRARPVSLTVPSQAGAQGRQFHGSASSLVEAVLISEGLGQFAQDPKFIEVTTQELADACDLTIEE 2104
CaV1.3 --SYRTFTPASLTVPSSFRNKNSDKQRSADSLVEAVLISEGLGRYARDPKFVSATKHEIADACDLTIDE 2092
CaV1.4 ---LRTFT--CLHVPGTHSDPSHGKRGSADSLVEAVLISEGLGLFARDPRFVALAKQEIADACRLTLDE 1924

                                  ICDI
CaV1.2 MENAADDILSGGARQSPNGTLLPFVNRRDPGRDRAGQNEQDASGACAPGCGQ-SEEALADRRAGVSSL* 2171
CaV1.3 MESAASTLLNGSVCPRANGDMGPISHRQDYELQDFGPGYSDEEPDPG-----REEEDLADEMICITTL* 2155
CaV1.4 MDNAASDLLAQGTS--------SLYSDEESILSRF------------------DEEDLGDEMACVHAL* 1966
```

**Figure S8:** Alignment of the DCT of rabbit $Ca_V1.2$, rat $Ca_V1.3$ and human $Ca_V1.4$ corresponding to the channels used in main text figures 1-3. Relevant regions as defined in the text are marked in blue. Previously reported PCRD and DCRD regions (1) are highlighted in pink; the known cleavage site for the DCT is indicated by the red arrow (1,2); previously identified phosphorylation site is outlined by a black box (3). Critical residues identified in this study are in bold.

| $Ca_V1.3$ IQ mutation | $K_{a,\ Ch}$ | $CDI_{max}$ |
|---|---|---|
| WT | 16.346 | 0.808 |
| F[4]A | 3.269 | 0.717 |
| F[-2]A | 22.231 | 0.747 |
| Y[-5]A | 7.192 | 0.802 |

**Table S1**: Parameters used to fit equation 1 to data in Fig. 1H. Values were originally measured in (4).
